# Experimental and natural peripheral HSV-1 infection: Neurotropism and impact on Alzheimer’s disease-related molecular markers

**DOI:** 10.64898/2026.05.07.723559

**Authors:** Axel Legrand, Susana Boluda, Marina Boukhvalova, Flore Rozenberg, Michel Bottlaender, Julien Lagarde, Marie Sarazin, Catherine Helmer, Morgane Linard, Benoît Delatour

**Affiliations:** Sorbonne Université, Institut du Cerveau - Paris Brain Institute - ICM, Inserm, CNRS, APHP, Hôpital de la Pitié Salpêtrière, Paris, France; Department of Neuropathology, AP-HP, Pitié-Salpêtrière Hospital, Paris, France; Sigmovir Biosystems, Inc., 9610 Medical Center Drive, Suite 100, Rockville, MD, 20850, USA; Department of Virology, Cochin University Hospital, Université Paris Descartes, 75014 Paris, France; Université Paris-Saclay, UNIACT, NeuroSpin, CEA, Gif-sur-Yvette, 91191, France; Université Paris-Saclay, BioMaps, Service Hospitalier Frédéric Joliot CEA, CNRS, Inserm, Orsay, France; Department of Neurology of Memory and Language, CMRR Sainte Anne Paris, GHU Paris Psychiatrie & Neurosciences, Hôpital Sainte-Anne, Paris, France; University of Bordeaux, INSERM, Bordeaux Population Health, UMR 1219, Bordeaux, France

## Abstract

Herpes Simplex virus type 1 (HSV-1) is a highly prevalent neurotropic virus from the alphaherpesviruses family. In recent years, a growing body of research has focused on the potential role of HSV-1 infections and recurrent reactivations in the pathophysiology of Alzheimer’s disease (AD). In particular, it has been hypothesized that HSV-1 could initiate or amplify the formation of neuropathological lesions characteristic of AD.

To explore further this hypothesis, we adopted an integrated approach aiming at deciphering the impact of HSV-1 infection on AD molecular markers (Aß and Tau pathologies) and combining experimental animal models of *in vivo* infection, postmortem neuropathological analysis of AD brains, as well as in-vivo clinical analysis in AD patients.

In animal models of peripheral (labial) infection with HSV-1 virus, we analyzed viral dissemination from peripheral tissues to the CNS, and the associated neuropathological consequences. Histological and molecular analyses revealed the occurrence of viral material (RNA, proteins) in the brainstem, the primary site of viral neuroinvasion, and in more anterior regions of the brain. Viral signatures were accompanied by early abnormal deposits of Aβ peptides and accumulation of phosphoTau (pTau) proteins in various brain areas.

Neuropathological examination of AD/control participants also underlined the presence of HSV-1 DNA in the human brainstem (pons) that was always associated with local Aß/Tau aggregates.

Finally, in AD patients, associations were found between HSV-1 seropositivity and neuropathological lesion burden (region-specific Tau and Aβ deposition detected by neuroimaging).

Taken together, these data provide new evidence in favor of the involvement of HSV-1 in the pathophysiology of AD, stressing a possible causal link between HSV-1 infection, neuroinvasion and AD neuropathological hallmarks (Aß lesions and tauopathy).

## Introduction

Alzheimer’s disease (AD) is currently the most common neurodegenerative disease, affecting millions of individuals worldwide (Nichols et al., 2022), and is characterized neuropathologically by extracellular deposits (plaques made of amyloid-ß peptides) and intraneuronal inclusions (neurofibrillary tangles constituted of hyperphosphorylated Tau proteins) (Duyckaerts et al., 2009). Despite significant advances in understanding the underlying mechanisms of AD, the exact causes of the disease remain unclear. Among the various etiopathological hypotheses explored, the infectious hypothesis has emerged as a particularly interesting avenue, supported by an increasing number of studies. The infectious hypothesis proposes that pathogens, particularly viruses, could play a role in the initiation and progression of AD neuropathology, therefore offering an alternative to current pathophysiological models (Itzhaki et al., 2020; Liu et al., 2025a; Zhao et al., 2024).

Various pathogens have been suspected, in particular the Herpes Simplex Virus Type 1 (HSV-1), a neurotropic virus that is highly prevalent in the population, whose infection persists throughout life (Steiner et al., 2007). HSV-1 infection occurs in three phases (Marcocci et al., 2020): the primary infection, often asymptomatic and usually occurring in childhood; the latent phase, during which the virus remains dormant in trigeminal ganglia (TG), no longer producing viral proteins but only the non-coding LAT RNA; and reactivation phases that occur throughout life, based on a complex balance between the host and the virus, and can be triggered by various stresses such as temperature, neurological trauma or immunosuppression. The TG are the main reservoir of latent viruses, and each ganglion is directly connected to the brainstem via the centripetal branch of its pseudo-bipolar neurons. Thus, the virus is theoretically able to propagate in the brainstem, after primo-infection or during reactivation episodes, and may reach specific subcortical nuclei, such as the locus coeruleus where early AD neuropathological changes (ADNC) have been described in human tissues (Braak et al., 2015; Grinberg et al., 2011; Theofilas et al., 2017). The neuroanatomical pathways connecting peripheral HSV-1 infection and CNS invasion have led to speculation that this virus may contribute to chronic local neuroinflammation and onset of AD neuropathology (Ball, 1982).

There are multiple evidences, mostly indirect, supporting an infectious hypothesis in AD and linking HSV-1 infection to dementia: 1) post-mortem studies have revealed the presence of HSV-1 viral DNA in human brain samples located within AD amyloid plaques (Wozniak et al., 2009b), 2) apolipoprotein E allele 4 (APOE4), a known major genetic risk factor in AD, is also a risk factor for HSV-1 infection (Itzhaki et al., 2006), 3) research has identified the beta-amyloid (Aß) peptide as an antimicrobial peptide, suggesting that amyloid deposits are part of an innate immune response to infection (Bourgade et al., 2016; Eimer et al., 2018; Gosztyla et al., 2018; Moir et al., 2018), 4) several epidemiological studies have also highlighted an increased risk of dementia in individuals with a history of HSV-1 infection and/or reactivations (Liu et al., 2025b; Lövheim et al., 2015; Tzeng et al., 2018) despite a lack of consistent analysis of the impact of HSV-1 infection on AD biomarkers (see however (Goldhardt et al., 2022)).

Moreover, experimental data also provided insights to reinforce a direct contributing role of HSV-1 in AD physiopathology. Both *in vitro* and *in vivo* evidence confirmed that HSV-1 infection could indeed initiate the accumulation of extracellular Aβ deposits and of neuronal Tau hyperphosphorylation (Wozniak et al., 2009a; Wozniak et al., 2007). In a recent study, De Chiara and colleagues (De Chiara et al., 2019) showed that peripherally HSV-1 infected wild-type mice develop cognitive impairments as well as abnormal Aß and phospho-Tau proteins (pTau) accumulation in the forebrain. AD-like induced phenotypes described in this latter study were not observed after primo-infection but required repeated rounds of viral reactivation to promote behavioral-neuropathological changes.

The present work aimed to fill several remaining gaps of the HSV-1/AD connection hypothesis: 1) what are the consequences of HSV-1 infection, in terms of ADNC, at the level of known early sites of neuroinvasion (brainstem); 2) could HSV-1 be detected, in humans, specifically at these primary sites of neuroinvasion ?; and 3) in AD patients, are there any associations between HSV-1 serostatus and AD-related pathophysiological biomarkers.

In order to address these points, we carried out infection experiments in cotton rats, a rodent species recurrently used to study human viral infections (Niewiesk et al., 2002) including HSV-1 (Boukhvalova et al., 2019; Lewandowski et al., 2002). Cotton rats, compared to regular inbred mice, are naturally more permissive to non-adapted human viruses and reproduce human-like tissue tropism, pathology, and immune responses more faithfully (Niewiesk et al., 2002). We also analyzed human postmortem brain samples (dorsal pons) from aged control and AD participants by HSV-1 PCR. Finally, clinical analysis was performed in a cohort of AD patients with available anti-HSV-1 serologies as well as MRI measures (to assess the integrity of the locus coeruleus, one of the first region showing ADNC (Ehrenberg et al., 2017)), amyloid and Tau PET molecular imaging and Aß/Tau cerebrospinal fluid biomarkers.

## Material & Methods

### Animal studies

#### Animals

Inbred *Sigmodon hispidus* cotton rats originated from a dedicated colony maintained by Sigmovir Biosystems, Inc., a private company that was used as subcontractor for the *in vivo* part of this preclinical study. The colony was monitored for the presence of antibodies to common rodent adventitious respiratory viruses and other pathogens, and no such antibodies were detected. Animals were housed in large polycarbonate cages and provided with standard rodent chow and water *ad libitum*. A total of 34 male and female cotton rats ranging from 5 to 6 weeks of age were used. All experimental procedures involving animals were conducted in accordance with applicable laws and guidelines and received mandatory approval from the Institutional Animal Care and Use Committee (IACUC) of Sigmovir Biosystems, Inc. Following lip inoculation with HSV-1 or PBS, rats were sacrificed to perform viral titrations in the lip and in PNS/CNS brain regions (infected rats) as well as neuropathological analysis and biochemical dosages in different brain regions of HSV1-infected and sham-infected rats (see table 1).

**Table 1.**
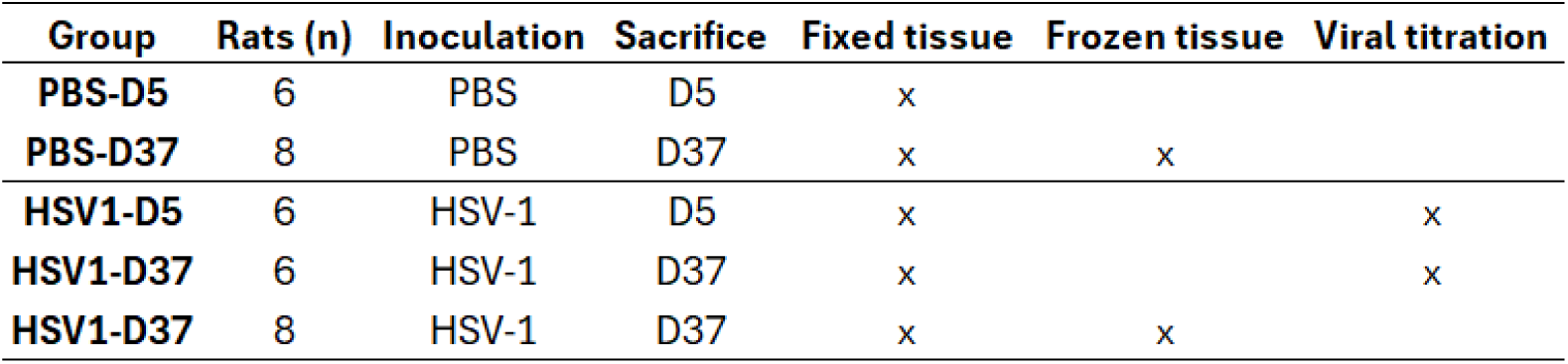
Animals and tissue processing. Rats were either lip-inoculated with HSV-1 or PBS and sacrificed 5 (D5) or 37 (D37) days later. Brain and peripheral tissue were collected and used for neuropathological/biochemical analysis or for titration of HSV-1 lytic activity.

#### Infection procedure

The protocol was adapted from (Boukhvalova et al., 2019). HSV-1 (strain 17) was propagated on Vero cells and cryopreserved at −80°C. Prior to inoculation, the virus was diluted to the target concentration in phosphate-buffered saline (PBS, pH 7.4), prepared within one hour of administration, and maintained on ice. Before inoculation, cotton rats were anesthetized via intramuscular injection of a ketamine-xylazine mixture (5 parts of 100 mg/ml ketamine and 1 part of 33 mg/ml xylazine, administered at a volume of 100 µl/100 g of body weight).

Inoculation was performed by (superficial) lip abrasion and involved placing a 20 µl drop of the virus solution containing 5 log_10_ PFU onto the vermilion border of the upper lip, bilaterally. Infection was achieved by subsequently performing repeated abrasion of the area using a sharp needle. Control animals for abrasion studies received 20 µl of PBS (pH 7.4) via the same abrasion method. All rats were carefully examined during the post-infection period to detect and monitor clinical signs.

#### Tissue harvesting

Animals were sacrificed by CO_2_ asphyxiation at 5 days (D5) or at 37 days (D37) after infection. Rats were either used for post-infection viral titrations (see below) or for histopathological and biochemical assays.

Hemibrains dedicated to neuropathological studies were fixed by immersion in formalin (4 days at RT) and then transferred to PBS until processing. Hemibrains used for subsequent ELISA dosage were macro-dissected in two rostro-caudal blocks: (i) posterior block comprising brainstem, cerebellum, and (ii) anterior block with the forebrain, and immediately snap-frozen and kept at-80°C until use.

#### Viral titration (plaques assay)

Tissue samples collected for virus quantification included the upper lip, the TG, and macro-dissected brain regions (brainstem, cerebellum, forebrain). The presence of infectious HSV-1 in tissue homogenates was determined by plaque assay on confluent Vero cell monolayers. Tissue samples were homogenized in EMEM medium containing sucrose stabilizing solution (homogenization buffer). Homogenates were then clarified by centrifugation and serially diluted in EMEM medium.

Diluted homogenates were used to infect Vero monolayers in 24-well plates, in duplicates. Following a 1-hour incubation period at 37°C in a 5% CO2 incubator, the cells were overlaid with medium containing 0.75% methylcellulose. After two days of incubation, the overlay was removed, and the cells were fixed and stained with 0.1% crystal violet solution for one hour. The plates were subsequently rinsed and air dried, and the resulting plaques forming units (PFU) were counted.

Viral titer was expressed as PFU per ml of inoculated solution for the lip, TG and brain, normalized according to the dilution factor used.

#### Histological analysis

Fixed hemibrains were sectioned, using a dedicated matrix (Young rat brain slicer matrix, Zivic Instruments, #BSRYS001-1) into 6 blocks encompassing the rostro-caudal extent of the brain, from the frontal pole to the brainstem and cerebellum (Rao et al., 2014).

Tissue blocks were then embedded in paraffin and cut with the help of a microtome (HM 340E, Thermo Scientific) to collect 4µm thick formalin-fixed paraffin-embedded serial sections (FFPE).

Standard hematoxylin and eosin (H&E) protocols were applied for general observation of tissue integrity, cell distribution and morphology. Immunohistochemistry (IHC) relied on immunoperoxydase-based methods with diaminobenzidine as chromogen. Primary antibodies targeted viral (HSV-1) proteins (anti-HSV-1 polyclonal #AB9533, Abcam, 1:1000; antigen retrieval: 30’ in citrate buffer at 90°C), Aß (anti-Aß 1-40/42 polyclonal #AB5076, Millipore, 1:750; antigen retrieval: 5’ in 98% formic acid), pTau (polyclonal anti-pTau205 #AB4841, Abcam, 1:500; antigen retrieval: 30’ in citrate buffer at 90°C). All antibodies were selected from previous publications on HSV-1 infected rodents (Boukhvalova et al., 2019; De Chiara et al., 2019) and protocols were optimized in positive control tissue before use.

*In situ* hybridization (ISH) of HSV-1 RNAs was also performed on FFPE sections using the RNAscope technology (ACD Bio-Techne) following manufacturer recommendations. Two probes were used to detect lytic RNA (RNAscope Probe-V-HSV-1-UL37-O1 #892001) or latent RNA (RNAscope Probe-HSV-1-LAT-C2 #315651-C2), as exemplified in previous work (Zhang et al., 2022).

All IHC/ISH sections were then digitized using a Nanozoomer scanner (Hamamatsu Photonics, Massy, France) at a 40X magnification to generate whole-slide images (WSI).

#### Biochemical analysis

Snap-frozen brain blocks were homogenized using two buffers: 1) HS-RAB buffer allowing extraction of highly soluble proteins, and 2) RIPA buffer permitting extraction of less soluble proteins such as membrane proteins or proteins in partially aggregated forms.

The tissues were first homogenized with a Dounce homogenizer in the HS-RAB buffer complemented with a cocktail of protease and phosphatase inhibitors (Pierce protease and phosphatase Inhibitor Mini Tablets, Thermo Fisher Scientific, #A32961). The homogenate obtained was subjected to sonication (Fisherbrand model 120 sonicator, 40 cycles of 1 second with an amplitude of 40%), then centrifuged at 7,000g at 4°C for 5 minutes in order to eliminate tissue debris. The collected supernatant was subsequently ultra-centrifuged at 100,000g at 4°C for 20 minutes. The final supernatant obtained corresponds to the HS-RAB fraction of the starting homogenate and was directly stored at-80°C. The pellet was resuspended and homogenized by pipetting in an equivalent volume of RIPA buffer, then centrifuged at 100,000g at 4°C for 20 minutes. The obtained supernatant corresponds to the RIPA fraction and was also stored at-80°C.

Total Tau and phospho-Tau (pT231) were quantified in duplicates using an MSD Kit (Phospho (Thr231) / Total Tau Kit, # K15121D; Meso Scale Discovery, Rockville, MD, USA) according to the manufacturer’s instructions. Signals were measured on a SECTOR Imager 2400 reader and analyzed with the Discovery Workbench software (Meso Scale Discovery).

### Human studies: postmortem analysis

#### Participants

Seventeen post-mortem brains, twelve confirmed AD cases (Braak V-VI, Thal 1-5) and 5 controls (Braak 0-III, Thal 0) with no clinical neurodegenerative pathologies, collected at Pitié-Salpêtrière hospital (Paris), were investigated (see table 4 in Results section). All brains were part of a brain donation project and stored in the French national brain biobank Neuro-CEB. One hemisphere was fixed in 4% formalin and the diagnostic of AD neuropathological changes included Aβ (6F3D, Dako) and Tau (AT8, Thermo Fisher Scientific) immunohistochemistry performed following routine protocols based on automated slide staining systems (Ventana Medical System Inc., Roche). The other hemibrain was snap-frozen and stored at-80°C for subsequent biochemical/molecular analyses requiring unfixed tissue.

#### Neuropathological examination and HSV-1 PCR

For each participant, brainstem FFPE sections at the level of the 4^th^ ventricle (rostral pons) were examined using H&E staining and anti-Tau/Aß immunostaining. This anatomical level encompasses both the locus coeruleus (LC) that is early affected during AD and part of the spinal trigeminal nucleus acting as a first relay for HSV-1 CNS neuroinvasion. Neuronal loss (easily evaluated by identifying the density of pigmented neurons in the area of the locus coeruleus) and Aß and Tau burden were semi-quantitatively scored from absence of lesions to florid pathology (0: lack, 1: weak, 2: moderate, 3: high).

For the 17 participants studied, tissue blocks from the frozen contralateral brain hemisphere were harvested in the same region of interest (pons tegmentum), i.e. in the vicinity of early sites of HSV-1 putative neuroinvasion. After tissue digestion and DNA extraction, PCR was performed to detect the presence/absence of HSV-1 DNA in the samples using reference protocols (Hausfater et al., 2004).

### Human studies: clinical analysis

#### Design and participants

We included 35 AD cases and 14 healthy elderly controls from the ShaTau7-Imatau cohort (https://clinicaltrials.gov/study/NCT0257682) (Table 2). AD patients were recruited according to the following criteria: (i) AD clinical phenotype; (ii) the positivity of AD pathophysiological biomarkers including both cerebrospinal fluid (CSF) biomarkers and amyloid PET imaging; (iii) Clinical Dementia Rating (CDR) scale ≤ 2. AD patients were classified as typical AD (predominant limbic/amnesic phenotype, n=20) and atypical AD (predominant cortical phenotype with relative preservation of memory function, n=15).

**Table 2.** Participants characteristics in the ShaTau subcohort. Numeric data are expressed as mean ± standard error to the mean. Metrics of AD cases and of typical-atypical AD variants are respectively in black and gray colors. CDR: clinical dementia rating scale; MMSE: Mini Mental State Examination, M(en), W(omen).

|  | Control (n=14) | AD (n=35) | typical AD (n=20) | atypical AD (n=15) |
| --- | --- | --- | --- | --- |
| Sex | 9W-5M | 21W – 14M | 12W – 8M | 9W – 6M |
| Age (years) | 68.4 ± 3.8 | 67.8 ± 7.9 | 70.5 ± 7.2 | 64.2 ± 7.6 |
| Disease duration (years) | - | 4.7 ± 3.3 | 5.25 ± 4 | 4 ± 2 |
| Education level (years) | 14.7 ± 3.3 | 14.1 ± 4.4 | 14.2 ± 4.2 | 14 ± 4.9 |
| MMSE | 28.8 ± 1 | 22.1 ± 4.8 | 24.2 ± 3.2 | 19.3 ± 5.4 |
| CDR | 0 | 0.5 (n=24) ; 1 (n=9) ; 2 (n=2) | 0.5 (n=16) ; 1 (n=4) | 0.5 (n=8) ; 1 (n=5) ; 2 (n=2) |
| Presence of APOE-ε4 allele | 2 | 22 | 14 | 8 |

Healthy elderly controls were included according to the following criteria: (i) Mini-Mental State Examination (MMSE) score ≥ 27/30; (ii) normal neuropsychological assessment; (iii) CDR = 0; (iv) no memory complaint; and (v) negative PET imaging. Exclusion criteria were previously described elsewhere (Lagarde et al., 2022).

All participants underwent the same procedure including a neuropsychological assessment, 3T brain MRI, ^11^C-PiB and ^18^F- Flortaucipir PET imaging. Blood samples were also drawn to determine the APOE genotype. For AD patients, the lumbar puncture was performed during routine clinical care before inclusion and CSF samples collected for research.

The Ethics Committee (Ile-de-France VI) approved the study. All subjects provided written informed consent.

#### Anti-HSV-1 serology

For all studied participants, frozen plasma samples taken at baseline were used for measuring anti-HSV-1 IgG. Antibody titrations were performed by routine procedures at the virology dpt. of Cochin Hospital (Paris) and relied on standard chemiluminescent ELISA with dedicated kit (LIAISON HSV-1 IgG, #310830, DiaSorin) and the LIAISON® XL fully automated, batch-processing analytical system (Diasorin). As recommended by the manufacturers, i) an index value ≥1.10 was considered to indicate the presence of IgG, ii) an index value < 0.9 was considered to indicate the absence of IgG and iii) no conclusion could be made for index values between 0.9 and 1.10. The used ELISA kit to detect IgGs is highly specific to HSV-1 and does not cross-detect, for instance, with anti-HSV-2 IgGs.

Statistical analyses were performed considering seropositive or seronegative status; the study of anti-HSV-1 IgG levels was limited by the small sample size and was not taken into account in multivariate regression analysis.

#### CSF AD biomarkers

CSF biomarkers including Aβ42, total Tau (t-Tau) and phosphorylated Tau (p-Tau181) were measured using ELISA with INNOTEST assays (Fujirebio, Ghent, Belgium).

#### MRI and PET imaging

All participants underwent a 3T MRI (Siemens PRISMA) which was performed at the Centre de Neuro-Imagerie de Recherche (Paris) including (a) three-dimensional (3D) T1-weighted volumetric magnetization-prepared rapid gradient echo (MP-RAGE) sequence, (b) neuromelanin-sensitive images acquired using two-dimensional axial turbo spin echo T1-weighted images that were positioned on the sagittal 3D T1-weighted images perpendicular to the posterior border of the brainstem from the lower part of the pons to the upper part of the midbrain, covering the entire LC. The analysis of neuromelanin-sensitive images was performed using a previously described automated method (Garcia-Lorenzo et al., 2013; Olivieri et al., 2019) in order to determine the signal intensity of the 10 brightest connected voxels bilaterally within the LC area, which was considered a reflection of the integrity/alteration of the LC.

Participants also underwent [^11^C]-PiB and [^18^F]-Flortaucipir PET at Service Hospitalier Frédéric Joliot (Orsay, CEA) on a High-Resolution Research Tomograph (HRRT; CTI/Siemens Molecular Imaging). PET acquisitions were performed at least 40 to 60 min after injection of 322.1 ± 68 MBq of [^11^C]-PiB to detect Aß deposits, and 80–100 min after injection of 376.8 ± 30.3 MBq of [^18^F]-Flortaucipir for the in-vivo labeling of Tau lesions.

Dynamic list mode acquisitions were binned into successive 5-min time frames. Post-processing image corrections were performed as previously described (Sureau et al., 2008; Varrone et al., 2009).

Parametric images were then created using BrainVisa software (http://brainvisa.info) on averaged images over 40–60min after injection of [^11^C]-PiB and over 80–100 min after injection of [^18^F]-Flortaucipir. Standard Uptake Value Ratio (SUVR) parametric images were obtained by dividing each voxel by the corresponding value found in the cerebellar gray matter eroded (4 mm) in order to avoid including the superior part of the cerebellar vermis, which is a site of Flortaucipir off-target binding, and to avoid partial volume effects from the CSF or occipital cortex. An automated segmentation of the gray matter was performed on the 3D T1 MR images of each subject using SPM8 software (Institute of Neurology, London, UK; http://www.fil.ion.ucl.ac.uk/spm/). Automatic segmentation of cortical structures using the Automated Anatomic Labeling (AAL) Atlas defined volumes of interest (VOI), which were applied on PET space in each subject after coregistration using a standard mutual information algorithm. The VOIs defined separately for the left and right hemispheres were pooled.

We first assessed the global cortical Aß and Tau SUVRs (global cortical index, GCI) in all AD cases, in order to obtain a general evaluation of lesion loads in each participant.

Then, in order to test the hypothesis of a predominant temporal vulnerability related to known HSV-1 neurotopism and to reduce between-patient heterogeneity in global burden due to disease evolution, we adjusted each regional cortical SUVR to the GCIs. For amyloid load, we defined an Aß-temporal index by the ratio: Aß SUVR in the inferior temporal cortex / Aß GCI (we chose the inferior temporal region because it is prone to accumulate early Aß deposits in AD (Grothe et al., 2015)). For Tau load, we defined (i) a Tau-temporal index by the ratio: tau SUVR in the temporal cortex / Tau GCI, and (ii) a Tau-temporo-limbic index by the ratio: Tau SUVR in a temporo-limbic meta-region / Tau GCI. The temporo-limbic meta-region was derived from (Jack et al., 2017) and comprises amygdala, parahippocampal, fusiform, lateral temporal and medial temporal cortices. For cortical regions supposed to be less prone to HSV-1 neuroinvasion (i.e. extra-temporal cortices), we defined a lateral and a medial (i.e. precuneus) parietal index for both Aß and Tau by calculating the following variables (i) lateral parietal Aß index by the ratio: Aß SUVR in the lateral parietal cortex / Aß GCI, (ii) lateral parietal Tau index by the ratio: Tau SUVR in the lateral parietal cortex / Tau GCI, (iii) Precuneus Aß index by the ratio: Aß SUVR in the precuneus / Aß GCI, (iv) Precuneus Tau index by the ratio: Tau SUVR in the precuneus / Tau GCI.

#### Statistical analysis

First, multivariate logistic regression models were performed to compare HSV-1 serostatus between AD cases and controls adjusting for age, sex and educational level. No adjustments were made for APOE4 status since only two controls were APOE4 positive. Second, in AD cases, multivariate linear regression models were performed to assess associations between HSV-1 seropositivity and i) AD biomarkers in the CSF, ii) LC signal intensity, iii) Tau and amyloid burden measured by PET. These analyses were adjusted for possible confounding factors (age, sex and presence of at least one APOE4 allele). Analyses were first performed in the whole AD group. Then, because typical amnesic and atypical AD could differ in terms of the chronology of the pathophysiological events, we conducted analysis in the two AD sub-groups separately. Normality of residuals and homoscedasticity were assessed using a histogram of residuals, a normal quantile-quantile plot of residuals, and a plot of residuals versus predicted values.

All statistical tests were two-tailed, and the threshold for statistical significance was 5%. Analyses were performed with SAS software (version 9.4, SAS Institute).

## Results

### Neuropathological consequences of experimental HSV-1 infection in the cotton rat model

#### Neurotropism of HSV-1

##### Viral titrations

In cotton rats euthanized 5 days post-infection (D5) (Figure 1A), high viral titers were detected in the lip, with a mean of about 15,000 PFU/mL. Substantial viral loads were also measured in the trigeminal ganglia (∼4,500 PFU/mL) and in the brainstem (∼3,600 PFU/mL), underlining spread of the virus from the inoculation site to the central nervous system via retrograde axonal transport along trigeminal afferents. Lower but still detectable levels of infectious viral particles were detected in the cerebellum (∼120 PFU/mL). In the forebrain, viral titers were close to the detection threshold (∼21 PFU/mL). Overall, results at D5 indicated a limited but measurable cerebral dissemination and propagation of HSV-1.

**Figure 1.**
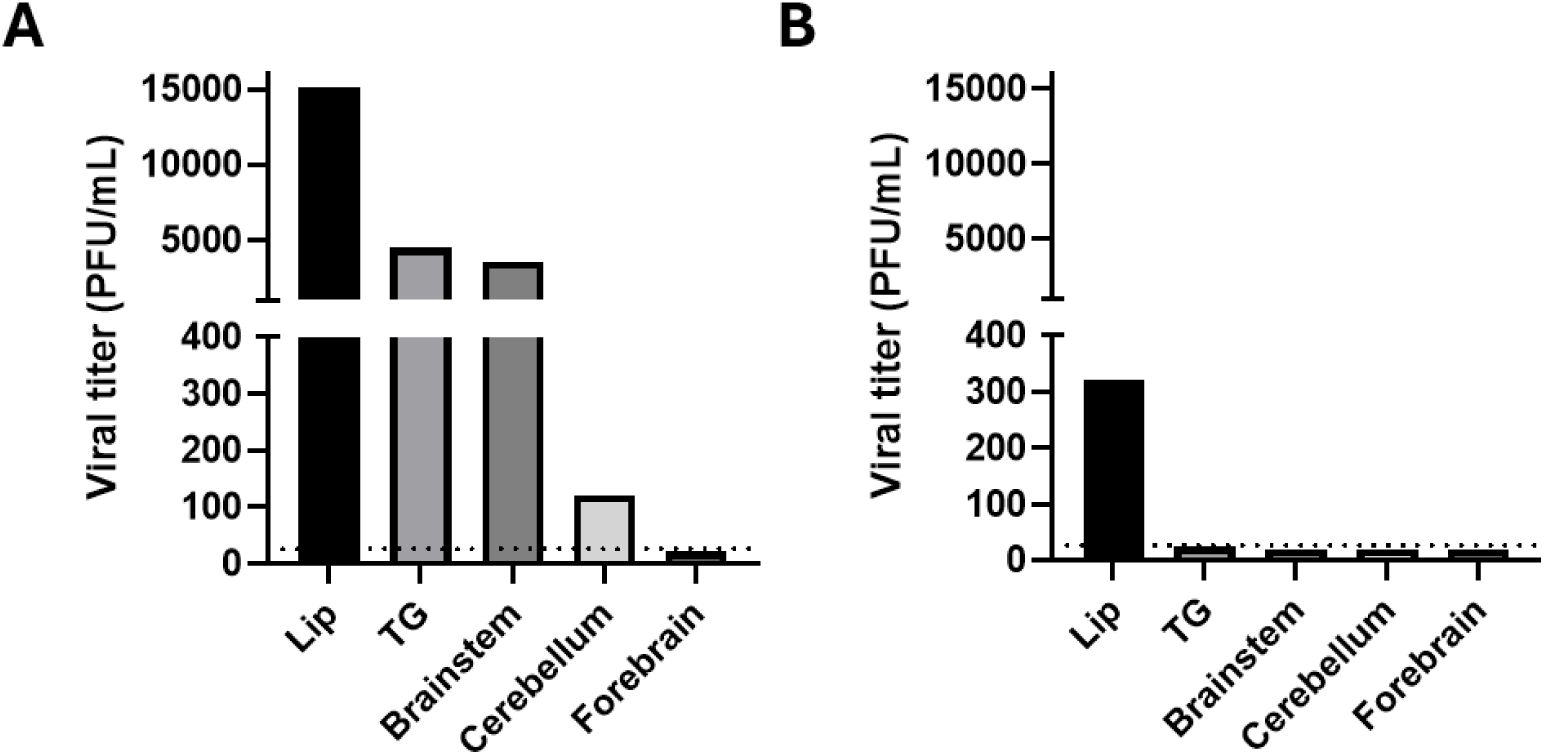
HSV-1 viral titrations at D5 and D37 The number of plaques forming units (PFU) was assessed in cotton rats sacrificed at D5 (A) or D37 (B) in peripheral tissue and in various PNS/CNS regions. Data expressed as geometric means.

At 37 days post-infection (D37), measured viral titers were markedly reduced across all analyzed tissues (Figure 1B). The inoculation site, the lip, still showed a low viral load (∼320 PFU/mL), while the trigeminal ganglia and CNS brain regions display values close to or at the limit of detection. This observation is consistent with the establishment of a latent phase in the sensory neurons of the trigeminal ganglia and with the absence of lytic virus in the brain.

##### General neuropathology and viral proteins immunohistochemistry

Gross histopathological analysis (H&E) staining revealed, as expected, characteristic brainstem and cerebellar tissue lesions in animals infected with HSV-1 and euthanized 5 or 37 days post-infection. These lesions were absent in uninfected animals (PBS groups). Lesions in HSV-1 infected rats consisted of inflammatory foci within the parenchyma (dense infiltrates of immune cells), perivascular cuffs, meningitis and areas showing vacuolated cells with perinuclear clearing, associated with an inflammatory infiltrate and/or the presence of glial cells, indicating an early stage of focal necrosis and neuronal degeneration. These lesions highlighted a marked neuroinflammatory process in the brainstem and cerebellum of infected cotton rats, which was consistent with HSV-1 neuroinvasion as already described in previous studies.

In D5 rats, viral proteins were immunodetected in the brainstem and in the cerebellar white matter (Figure 2.A,B). In contrast, no immunoreactivity could be observed in forebrain structures, which suggests, at this stage, a lack of viral spread to anterior brain regions in the cotton rat model. In D37 animals, the presence of HSV-1 viral proteins was not detected in any brain region analyzed (Figure 2.C,D). This observation could reflect a clearance of viral antigens 37 days post-infection and is compatible with a status of quiescence (HSV-1 latent phase with no active viral replication).

**Figure 2.**
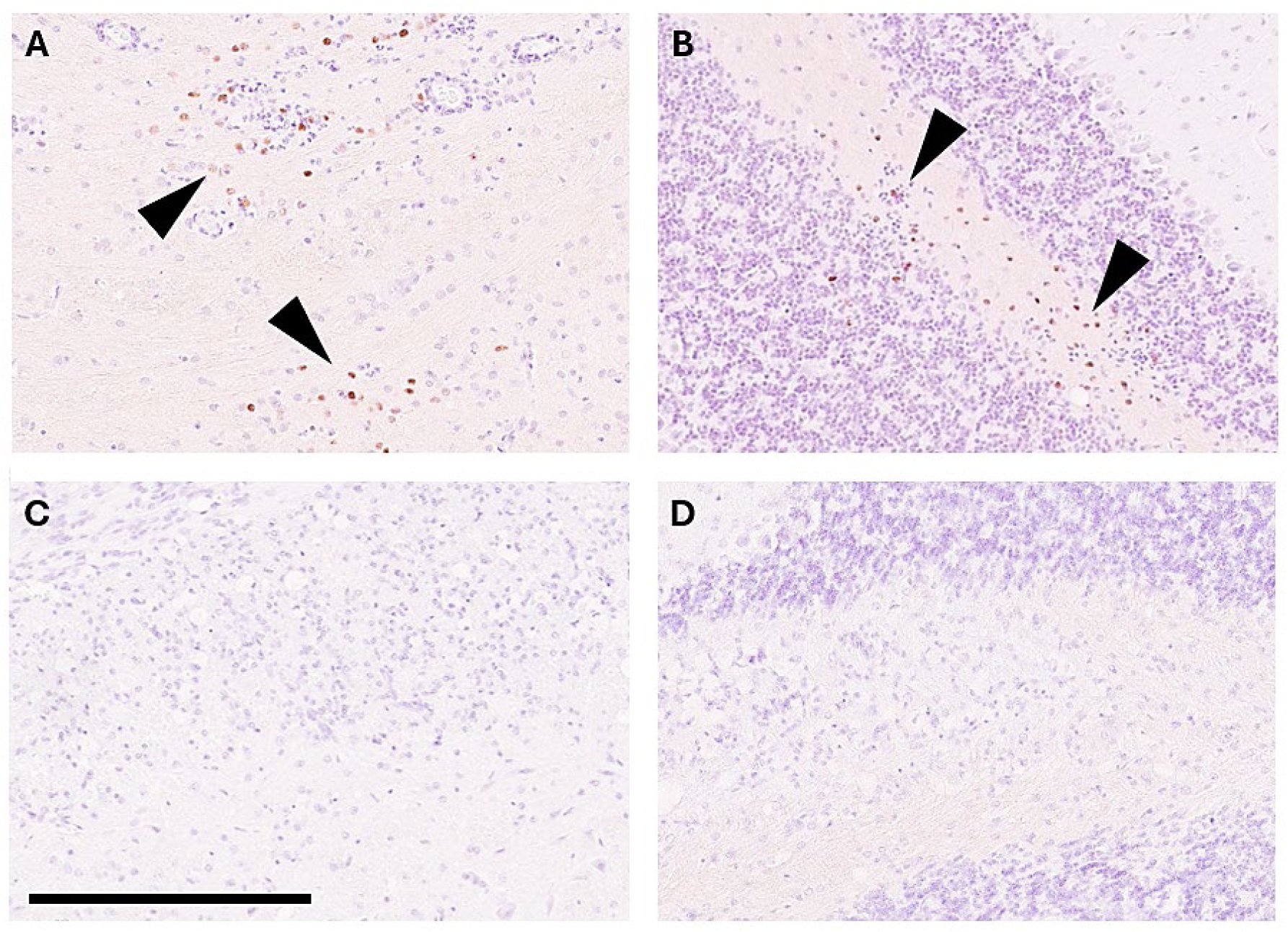
Viral proteins at D5 and D37 IHC was performed to detect viral proteins in rats sacrificed at D5 (A: brainstem, B: cerebellum white matter) or at D37 (C: brainstem, D: cerebellum white matter). Black arrow heads point to foci of positive immunodetected material. Scale bar 250 µm

##### Viral genome

In cotton rats euthanized at D5, both viral UL37 (lytic phase) and LAT RNAs were detected exclusively in the brainstem and cerebellum (Figure 3.A-B) with no presence of viral genome in other brain regions. The signals obtained were mostly restricted to a few localized foci, without widespread dissemination in the brain parenchyma. The data indicate that, even at early stages post-infection, lytic and quiescent HSV-1 genomes are present in the primary sites of CNS neuroinvasion.

**Figure 3.**
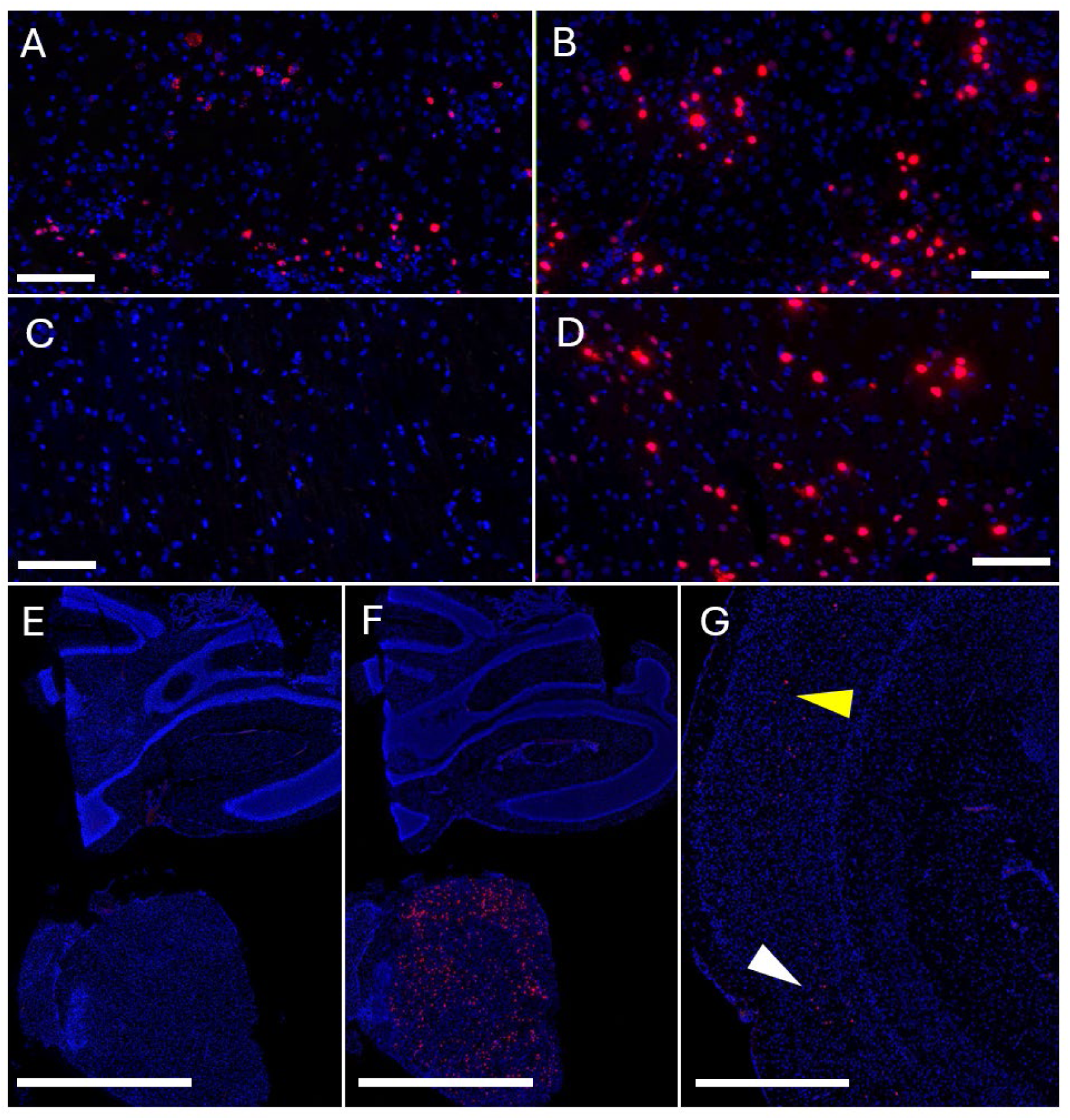
Viral RNAs at D5 and D37 (previous page) Lytic and latent HSV-1 RNAs were detected by RNAscope ISH in brain tissue from infected rats sacrificed at D5 (A,B) or D37 (C-G) post-infection. A. D5, lytic RNA, brainstem B. D5, latent RNA, brainstem C. D37, lytic RNA, brainstem D. D37, latent RNA, brainstem E. D37, lytic RNA, whole section with brainstem and cerebellum F. D37, latent RNA, adjacent whole section with brainstem and cerebellum G. D37, latent RNA in the isocortex. White arrowhead points to foci in the temporal (entorhinal-perirhinal) cortex while yellow arrowhead points to foci in the visual cortex. Scale bars: 250 µm (A-D), 2.5 mm (E-F), 1 mm (G)

**Figure 4.**
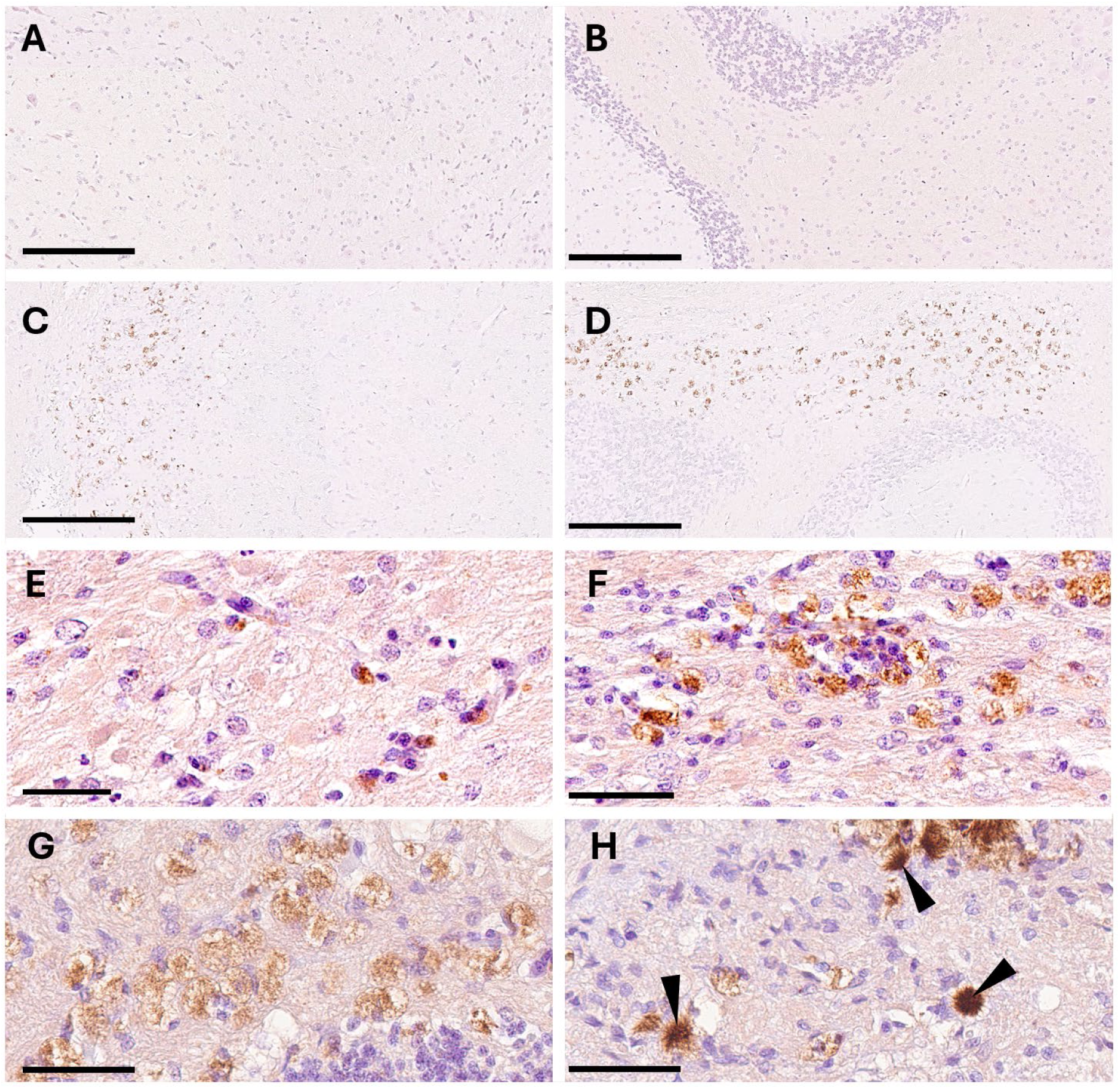
Aß deposits in HSV-1 infected rats. The presence of abnormal Aß deposition was scrutinized in infected rats sacrificed at D5 (A-B) and D37 (C-H). Rats sacrified at D5 (A-B) showed absence of Aß deposition, while in rats sacrificed at D37 (C-H) either intracellular(C-G) or extracellular fibrillar (H) immunoreactivity was seen. A. D5-brainstem B. D5-cerebellum C. D37-brainstem D. D37-cerebellum E. D37-thalamus F. D37-corpus callosum G. D37-intracellular immunoreactivity (brainstem) H. D37-extracellular immunoreactivity (brainstem, arrow heads pointing to extracellular deposits) Scale bars: 250 µm (A-D), 50 µm (E-H).

At D37 post-infection, the distribution pattern of HSV1 RNA transcripts was strikingly different: there was a lack of UL37 RNA (Figure 3.C,E) and an extensive and not topographically organized distribution of LAT transcripts, mainly in the brainstem (Figure 3.D,F) and, to a lesser extent, in more anterior brain structures of the diencephalon (eg thalamus) and isocortex (Figure 3.G).

Rats infected with PBS did not show, as expected, any specific labeling and background staining was mainly associated with blood vessels and easily identified.

#### Induction of AD-like proteinopathies

##### Amyloid-ß peptides deposition

Immunohistochemical analysis of the brains of cotton rats infected with HSV-1 did not reveal the presence of amyloid in animals euthanized at D5 post-infection (n=6 analyzed rats; Figure 3.A,B). However, immunohistochemical analysis performed on D37 animals (n=13) revealed the presence of Aβ-positive deposits in the brainstem and cerebellum (Figure 3.C,D), as well as in more anterior regions, including the thalamus (Figure 3.E) and the corpus callosum (Figure 3.F). Aß-positive deposits were more frequent in the posterior brain areas (69% of rats presenting Aß+ deposits) than in the forebrain regions (38% of rats being Aß+). Detected Aß deposits displayed two distinct features: 1) intracellular labeling, localized within vacuolar structures, suggesting an accumulation of Aβ peptides within phagocytic macrophage-like cells (Figure 3.G), and 2) extracellular labeling, characterized by compact, fibrillar-appearing deposits (Figure 3.H). Both types of deposits were found indiscriminately across the brain regions analyzed. No Aß labeling was noted in PBS-inoculated rats.

##### Tau hyperphosphorylation

A total of 22 animals were processed (6 PBS-treated rats sacrificed at D37, 3 D5-HSV-1 rats and 13 D37-HSV-1 rats) through anti-pT205 immunostaining.

Analysis showed that, in the cerebellum, an endogenous low-level of Tau phosphorylation was present in PBS-inoculated rats (Figure 5.A). An elevation of the overall pTau T205 signal was detected in the brains of infected cotton rats and the increase in Tau phosphorylation was stronger in animals euthanized at D37 (Figure 5.E) than in those euthanized at D5 (Figure 5.C). This overall signal elevation was predominantly observed in the white matter.

**Figure 5.**
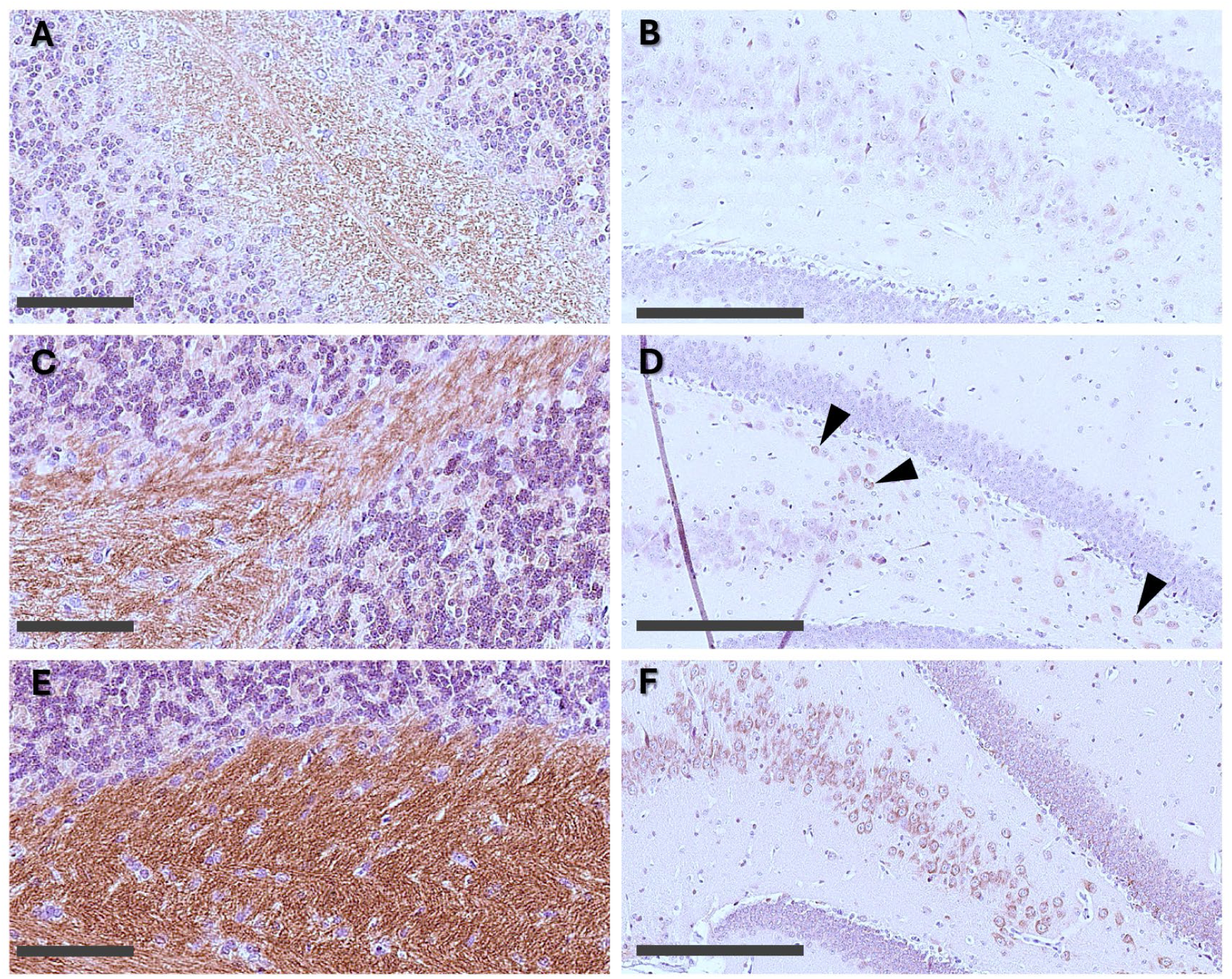
Tau abnormal phosphorylation (T205) in HSV-1 infected rats. Immunodetection of phospho-Tau at the T205 epitope was performed in PBS-uninfected rats (A,B) and in D5 (C,D) and D37 (E,F) infected rats. A.PBS-cerebellum white matter B.PBS-hippocampus CA3, hilus of dentate gyrus C.D5 HSV-1-cerebellum white matter D. D5 HSV-1- hippocampus CA3, hilus of dentate gyrus; arrowheads pointing at positive cells. E. D37 HSV-1-cerebellum white matter F. D37 HSV-1- hippocampus CA3, hilus of dentate gyrus. Scale bars: 100 µm (A,C,E); 250 µm (B,D,F)

**Figure 6.**
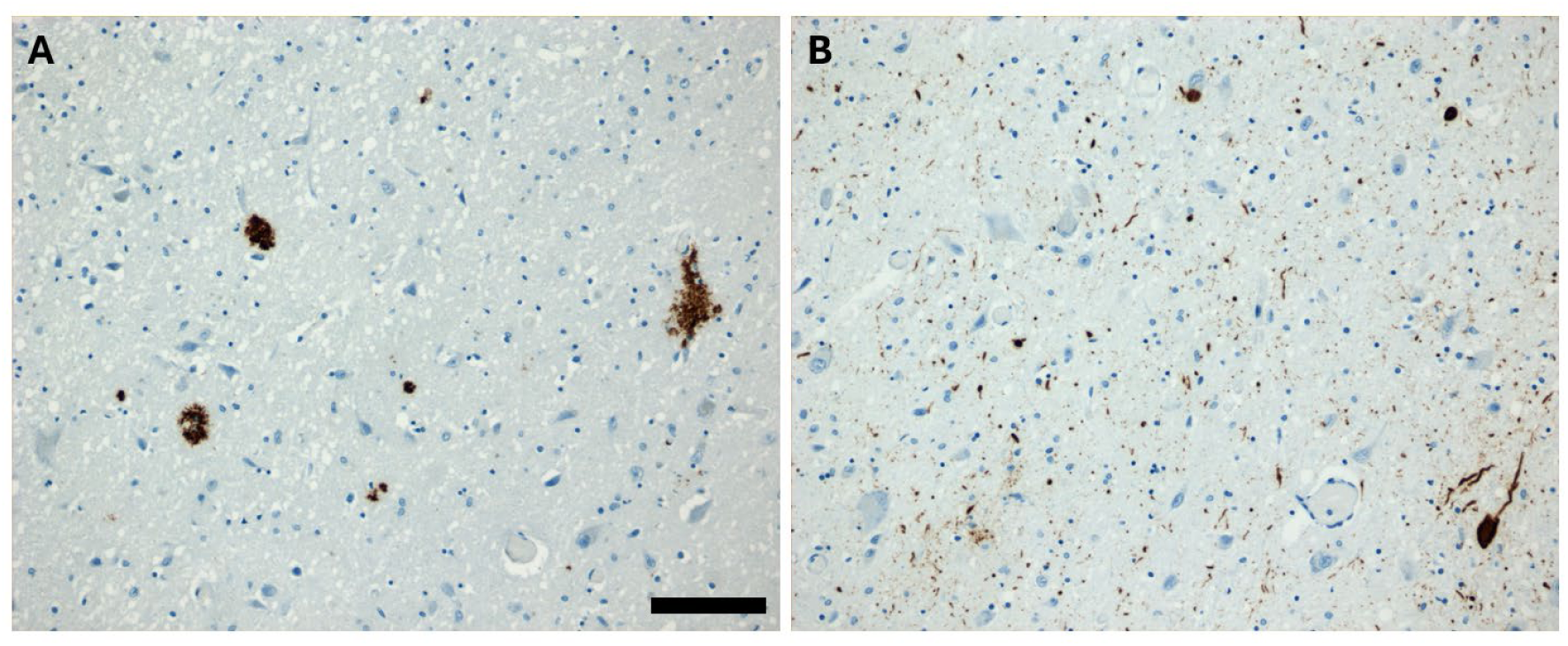
Alzheimer’s disease neuropathological changes observed in AD brainstem (dorsal pons). All examined AD cases displayed Aß deposits (A) and tauopathies (B) in the dorsal pons. The density of lesions was variable between cases. With limited exceptions (see Table 4) lesions were absent in control brains. Scale bar: 50 µm

In the forebrain, at 5 days post-infection, scarce pTau T205 neuronal immunoreactivity was observed in the hippocampus, specifically in the hilus of the dentate gyrus as well as in the CA3 pyramidal cell layer (Figure 5.D), in comparison to uninfected animals (Figure 5.B). In D37 animals, a stronger increase in pT205 immunoreactivity was noted in the same regions (Figure 5.F).

It appears therefore that constitutive endogenous Tau phosphorylation is abnormally increased after HSV-1 infection, both in the cerebellum and in the forebrain, and that Tau phosphorylation is more prominent at D37 than shortly after infection at D5.

In addition to histological observations, we measured pTau (T231) and total Tau in the brains of these cotton rats using MSD technology. The data obtained showed, for the RIPA fraction, a significant increase in the pTau / total Tau ratio in the forebrain of D37 rats compared to uninfected animals (Table 3). The same trend was observed for the RAB fraction but did not reach statistical significance. In the posterior brain (brainstem and cerebellum), no significant difference in the pTau/Tau ratio was detected between infected and uninfected cotton rats, regardless of the fraction examined.

**Table 3.** pT231 MSD dosages in control and HSV-1 infected rats. Dosages are expressed as mean of ratio pTau/total Tau +/- SEM. P-values are related to the comparison of PBS and HSV-1 groups (unpaired t-test).

| Fraction | Brain region | PBS | HSV-1 | P-value |
| --- | --- | --- | --- | --- |
| RAB | Brainstem - Cerebellum | 0.153 ±0.04 | 0.1903 ±0.017 | ns |
|  | Forebrain | 0.1775 ±0.007 | 0.1943 ±0.004 | ns |
| RIPA | Brainstem - Cerebellum | 0.389 ±0.103 | 0.314 ±0.015 | ns |
|  | Forebrain | 0.198 ±0.001 | 0.223 ±0.001 | p<.001 |

**Table 4.** Neuropathological evaluation and HSV-1 DNA detection. All studied participants were Braak-and Thal-staged. A semi-quantitative evaluation of cell loss in the locus coeruleus (LC) and Aß/Tau immunoreactivities was performed on FFPE sections. All individuals were also tested with HSV-1 PCR on frozen tissue extracted from the contralateral pons region (neg: negative PCR; POS: positive PCR).

|  | Sex | Age | Braak | Thal | LC loss | Aβ | pTau | HSV1 PCR |
| --- | --- | --- | --- | --- | --- | --- | --- | --- |
| AD THAL 5 | M | 82 | Braak VI | Thal 5 | 2 | 2 | 3 | neg |
|  | F | 84 | Braak VI | Thal 5 | 1 | 2 | 2 | **POS** |
|  | F | 86 | Braak VI | Thal 5 | 2 | 1 | 1 | **POS** |
|  | M | 87 | Braak VI | Thal 5 | 2 | 2 | 2 | neg |
|  | F | 89 | Braak VI | Thal 5 | 0 | 2 | 2 | neg |
|  | M | 90 | Braak VI | Thal 5 | 2 | 1 | 2 | **POS** |
|  | F | 94 | Braak V | Thal 5 | 1 | 1 | 3 | neg |
| AD THAL 1-3 | F | 93 | Braak V | Thal 1-3 | 0 | 0 | 1 | neg |
|  | F | 101 | Braak V | Thal 1-3 | 0 | 0 | 1 | neg |
|  | M | 86 | Braak VI | Thal 1-3 | 2 | 2 | 2 | neg |
|  | F | 74 | Braak VI | Thal 1-3 | 1 | 0 | 3 | neg |
|  | M | 87 | Braak VI | Thal 1-3 | 0 | 0 | 2 | neg |
| CTRL | F | 52 | Braak 0 | Thal 0 | 0 | 0 | 0 | neg |
|  | F | 60 | Braak 0 | Thal 0 | 0 | 0 | 0 | neg |
|  | M | 69 | Braak 0 | Thal 0 | 0 | 0 | 0 | neg |
|  | F | 92 | Braak I | Thal 0 | 0 | 0 | 1 | **POS** |
|  | M | 70 | Braak III | Thal 0 | 0 | 0 | 1 | neg |

### HSV-1 infection in humans

#### Neuropathological study: neurotropism of HSV-1

All AD cases with postmortem examination did present abnormal Aß and Tau proteinopathies in the brainstem region of the dorsal pons (Figure 1, Table 4). Lesion burden varied across cases, with increased densities of Aß- and Tau-positive deposits/inclusions in patients diagnosed with high Thal staging (Thal 5 vs Thal 1-3). Neuronal loss in the locus coeruleus was also variable and found to be more important in cases with high grading (Thal 5). Controls were devoid of any neuropathological alterations, with the exception of discrete pTau inclusions (2/5 participants).

In 25% (3/12) of AD cases the genome of HSV-1 was detected by PCR in the dorsal pons. All the HSV-1 positive AD cases did belong to the most advanced neuropathological subgroup (Braak VI, Thal 5, LC cell loss). Also one control individual was identified as HSV-1 positive and this participant did harbor immunodetected Tau lesions.

#### Clinical study

##### HSV-1 seropositivity in AD patients and controls

The analysis of HSV-1 seroprevalence in the studied cohort highlighted, as expected, a high exposure frequency, above 60%, in agreement with epidemiological data from the general population. Despite a slightly higher seroprevalence in the AD group, there was no statistically significant difference between AD patients and controls (77% vs 64%; adjusted Odds Ratio (aOR) =1.7 95% confidence interval [0.4 – 7.5] p=0.47), even when considering typical and atypical AD separately. Analysis of anti-HSV-1 IgG levels, although limited by the small-sized population and by occurrence of out-of-range measures in a number of individuals, did not show obvious differences between groups (eg controls vs AD, typical AD vs atypical AD; data not shown).

##### Relationships between HSV-1 seropositivity and AD biomarkers

Among all AD cases, no associations were found between HSV-1 seropositivity and levels of CSF biomarkers (Aß42, log-transformed pTau, total Tau, ratio Aß42/pTau, ratio Aß42/Tau (Table 5). Also, no associations were found between HSV-1 seropositivity and the LC signal intensity.

**Table 5.** Multivariate linear regression models assessing association between HSV-1 seropositivity and AD biomarkers. CSF dosages, locus coeruleus (LC) signal intensity and global Aß or Tau SUVRs were analyzed in the whole AD cohort (n=31-35) while brain regional Aß/Tau SUVRs are presented only in typical AD participants (n=18-19). *Adjusted for age, sex and presence of at least one APOE4 allele. ^£^P-Tau levels were log-transformed for linear regression models. ^§^For Aß ratios, the SUVRs were reported relative to the overall SUVR of grey matter. For Tau ratios, the SUVRs were reported relative to the overall SUVR of the cortex.

| ALL AD CASES | Mean (std) | HSV-1 seropositive<br>(vs seronegative)*<br>$\beta$ (std); p |
| --- | --- | --- |
| CSF Aβ42 n=35 | 456 (130) | $\beta$ - 59 (55); p 0.29 |
| CSF Tau n=35 | 683 (413) | $\beta$ -266 (178); p 0.15 |
| CSF P-Tau<sup>§</sup> n=35 | 91 (42) | $\beta$ -0.19 (0.18); p 0.30 |
| LC signal intensity n=35 | 1.22 (0.04) | $\beta$ 0.01 (0.02); p 0.47 |
| Aβ global SUVR n=33 | 2.9 (0.6) | $\beta$ 0.01 (0.29); p 0.97 |
| Tau global SUVR n=31 | 2.2 (0.9) | $\beta$ -0.32 (0.31); p 0.31 |
| TYPICAL AD CASES | Mean (std) | HSV-1 seropositive<br>(vs seronegative)*<br>$\beta$ (std); p |
| Aβ temporal ratio<sup>§</sup> n=19 | 0.86 (0.05) | $\beta$ 0.07 (0.02); p 0.01 |
| Aβ parietal ratio<sup>§</sup> n=19 | 0.91 (0.06) | $\beta$ -0.02 (0.04); p 0.60 |
| Aβ precuneus ratio<sup>§</sup> n=19 | 1.06 (0.10) | $\beta$ 0.05 (0.07); p 0.55 |
| Tau temporal ratio<sup>§</sup> n=18 | 1.13 (0.09) | $\beta$ 0.14 (0.05); p 0.03 |
| Tau temporo-limbic ratio<sup>§</sup> n=18 | 1.16 (0.11) | $\beta$ 0.15 (0.08); p 0.08 |
| Tau parietal ratio<sup>§</sup> n=18 | 1.01 (0.12) | $\beta$ -0.15 (0.07); p 0.04 |
| Tau Precuneus ratio<sup>§</sup> n=18 | 1.06 (0.18) | $\beta$ -0.05 (std 0.13); p 0.69 |

In the whole group of AD, we found no relation between the Aß GCI or tau GCI and HSV-1 seropositivity. In typical amnesic AD, however, a positive association was underscored between HSV-1 seropositivity and the Aß-temporal index (Table 5, Figure 7). This association was not observed in extra-temporal regions (lateral parietal cortex, precuneus). A positive association was also evidenced between HSV-1 seropositivity and the temporal-tau index and a similar positive association was noted with the temporo-limbic tau index but did not reach statistical significance (Table 5, Figure 7).

**Figure 7.**
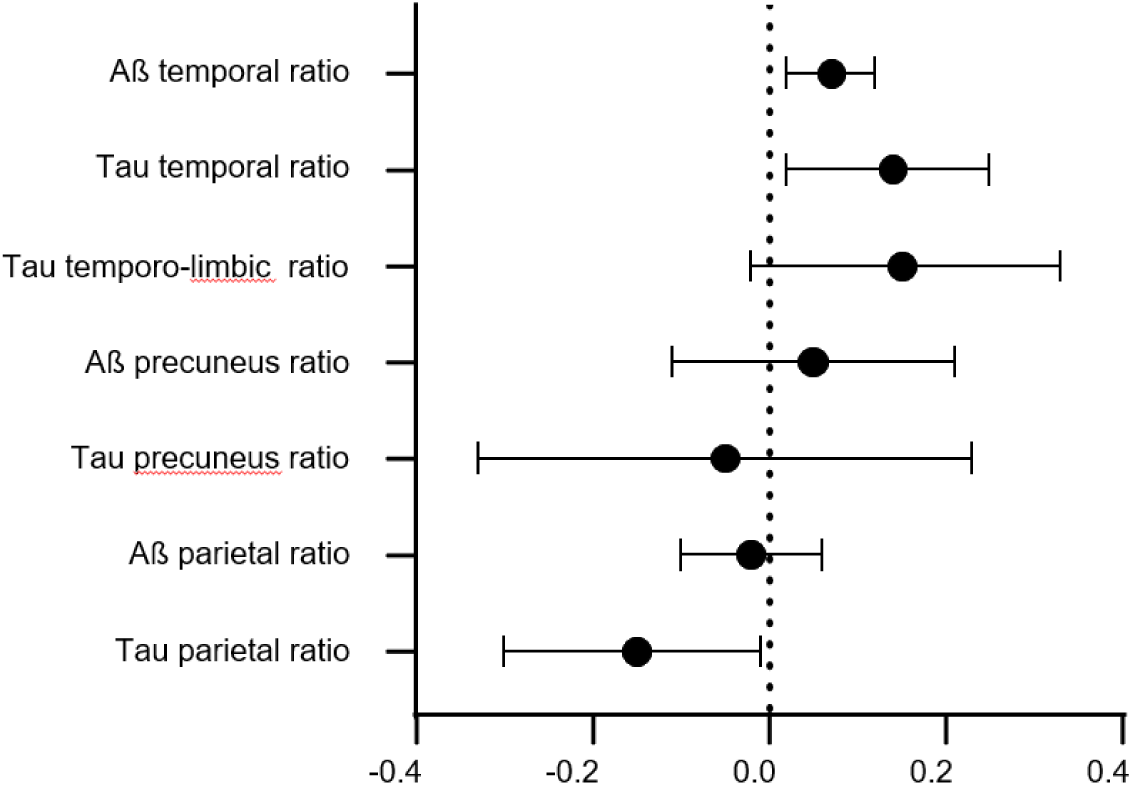
Association between HSV-1 seropositivity and Aß or Tau SUVRs in typical AD cases. Multivariate linear regression models adjusted for age, sex and APOE4 genotype were performed to evaluate association between HSV-1 seropositivity with temporal SUVR ratios (Aß temporal, Tau temporal, Tau temporo-limbic) or with extra-temporal SUVR ratios (Aß precuneus, Tau precuneus, Aß parietal, Tau parietal). For each graph, ß +/-95% confidence intervals are plotted (ß>0: positive association, ß<0: negative association).

Remarkably an opposite (negative) association was detected between HSV-1 seropositivity and the tau-parietal index in the typical AD subgroup.

In atypical AD patients, on the contrary, regression analysis did not underline any significant associations between HSV-1 seropositivity and Aß or tau regional indices (all ps>0.16). These analyses were, however, limited by a low statistical power due to the reduced number of atypical AD cases in the cohort.

## Discussion

HSV-1 is a human neurotropic alphaherpes virus that establishes a lifelong latent infection in sensory neurons. Following primary replication at mucosal surfaces (e.g., orofacial), the virus enters peripheral nerve endings and undergoes retrograde axonal transport to the cell bodies residing in TG. A critical, yet less frequent, outcome of infection is the invasion of the CNS, which can lead to herpes simplex limbic encephalitis. The impact of viral infection on the risk of developing neurodegenerative diseases has been repeatedly emphasized (Levine et al., 2023) and HSV-1, in particular, has been identified as a putative actor in the pathophysiology of AD (Harris et al., 2018). Still, the HSV-1/AD connection hypothesis relies mostly on indirect (epidemiological) and limited (experimental) support.

In the present study, we first confirmed, in the cotton rat model, the capacity of HSV-1 to neuroinvade the brainstem and distant brain areas following lip inoculation and to establish latency in these regions. More importantly, we evidenced that the brain penetration of HSV-1 was able to induce, after a single primary infection, local Aß abnormal accumulations in the brainstem/cerebellum, thalamus, and corpus callosum, as well as Tau hyperphosphorylation in cerebellum and downstream brain regions (hippocampus). The presence of HSV-1 was also evidenced in the pons of aged human individuals (either AD or controls), always associated with local Aß deposition and/or Tau aggregates. Finally, clinical analysis suggested an association between HSV-1 infection and Tau or Aß burden assessed by PET imaging in the temporal cortex in typical amnesic AD patients.

Animal data, primarily relying on murine models, have been instrumental in delineating the pathways, kinetics, and consequences of HSV-1 neuroinvasion. In cotton rats, we showed that peripheral (labial) HSV-1 infection led to brainstem dissemination of productive viruses at short term (D5: presence of viral proteins/genomes) followed by progressive quiescence (D37: absence of viral proteins, exclusive LAT RNA expression), in accordance with the previous data from the literature (Boukhvalova et al., 2022; Boukhvalova et al., 2019; Lewandowski et al., 2002). To our knowledge, a detailed characterization of lytic/latent HSV-1 RNAs in the brain has not been described in infected cotton rats. The present study indicates that while viral genome was strictly confined in the brainstem (primary sites of neuroinvasion) shortly after infection, a widespread distribution of LAT RNAs was observed during latency, suggesting progressive dissemination of HSV-1 from the brainstem to distant cortical and subcortical areas. Using the same ISH protocols as us (RNAscope technology), Zhang and collaborators (Zhang et al., 2022) also demonstrated the presence of HSV-1 RNAs in the brain of infected mice, in particular the co-expression of UL37 and LAT RNAs in the hippocampus at 5 weeks post-infection. These results cannot however be directly compared with our observations as a different route of inoculation that did not strictly mimic the natural pathways of HSV-1 infection (intravenous injection) was used by Zhang and colleagues.

Previous *in vitro* data have underlined the potential of HSV-1 to promote expression of APP fragments and Aß accumulation (De Chiara et al., 2010; Wozniak et al., 2007). *In vivo* inoculations in wild-type mice were occasionally reported to induce the formation of amyloid deposits in cortical areas (Wozniak et al., 2007), but using the intranasal route of infection. In young 5xFAD transgenic mice modeling AD brain amyloidosis, the intracranial injection of HSV-1 also rapidly promote the appearance of Aß plaques (Eimer et al., 2018), but these results were found to be non-reproducible a few years later (Bocharova et al., 2021). Conversely, few studies investigated labial infection (the principal, natural, and most common peripheral entry site for HSV-1 during primary infection in humans) and its consequences, in animal models, in terms of AD-like neuropathological changes. In wild-type BALB/c mice, labial infection was associated with isocortical-hippocampal accumulation of Aß peptides but only after several rounds of viral reactivation (De Chiara et al., 2019).

For the first time we evidenced that, following primo-infection using labial inoculation, abnormal Aß deposits were induced in the neuroinvaded areas of the brainstem and cerebellum, intermingled with neuroinflammatory territories. Aß deposits were found both intracellularly, presumably as microglia/macrophages phagocytocized material or alternatively as intraneuronal aggregates as described by others (Albaret et al., 2023), and extracellularly, with the appearance of fibrous round-shaped aggregates. These lesions were not observed at D5 but only at D37, suggesting a delayed reactive response. One could hypothesize that attempts to control viral neuroinvasion may first rely on neuroinflammatory regulations at short-term and later on, as a second line of defense, involve production of amyloid peptides known to trigger an innate immune anti-microbial action (Moir et al., 2018).

Similarly, previous *in vitro* data from the literature pinpointed the consequences of HSV-1 infection in promoting abnormal Tau hyperphosphorylation (Wozniak et al., 2009a).

Infected mice also develop AD-like tauopathies, after intranasal inoculation (Martin et al., 2014) or following repeated rounds of reactivation after labial inoculation (De Chiara et al., 2019).

Our *in vivo* data indicate that labial primo-infection in cotton-rats can induce abnormal Tau phosphorylation in the cerebellar white matter and also, at distance, in the hippocampus. As for Aß deposition, Tau anomalies were increased at D37, suggesting again a delayed response. Similar results were reported by Martin and colleagues (Martin et al., 2014) with an increase of pTau cortical immunoreactivity at D60 compared to D5 post-infection. Late Tau phosphorylation in cotton rats was particularly evidenced in anterior brain regions. Biochemical dosages indeed confirmed an elevation of the pTau/Tau ratio in the forebrain samples of D37 rats.

In our experiments, anti-Tau immunohistochemistry relied on the detection of the phosphorylated T205 epitope as exemplified in (De Chiara et al., 2019). Previous work showed that pT205 changes can be triggered by white matter anomalies (Strain et al., 2022) and one may suspect that HSV-1 induced demyelination (Boukhvalova et al., 2019) could contribute to Tau phosphorylation. Alternatively a role of pTau in censing damage-associated molecular patterns (DAMPs) as hypothesized by others (Powell-Doherty et al., 2020) remains also compatible with our observations.

The brainstem has for years been the poor relation of the research in AD, a pathology initially conceived as mainly targeting isocortical and archicortical (eg hippocampus) areas of the brain. By extension, most of previous work in postmortem brain tissue focused on the search for HSV-1 signatures in cortical regions and to our knowledge there was no report on the impact of HSV-1 neuroinvasion on AD neuropathological changes at the brainstem level. It is however considered that the mesencephalic and pons nuclei (upper and mid parts of the brainstem) are a source of interest and a critical hub at the early stages of AD (Grinberg et al., 2011; Rüb et al., 2016). AD neuropathological changes can indeed be detected in specific nuclei of the brainstem, such as the locus coeruleus or dorsal raphe nuclei but also the mesencephalic trigeminal nucleus that relays HSV-1 neuroinvasion (Klosen et al., 1993).

In the present work, the neuropathological examination of human brainstem (dorsal pons) from aged human individuals allowed, for the first time, to detect the local presence of HSV-1 DNA in multiple individuals. All subjects that were PCR-diagnosed as HSV-1 positive, including one aged non-demented individual, did harbor various amounts of Aß plaques and/or tauopathies. These results, although purely correlative and limited by the low number of examined participants and the variability and volume of tissue sampling, suggest that HSV-1 genome, when detected in the brainstem, is constantly accompanied by local AD-related neuropathological changes.

In the clinical cohort, we did not observe significant differences in terms of HSV-1 seroprevalence between AD and control participants, replicating some earlier observations (Lövheim et al., 2015). While no association were found between HSV-1 seropositivity and AD CSF biomarkers or LC signal intensity in AD cases, PET imaging data revealed discrete but significant associations between HSV-1 seropositivity and higher Aß and Tau temporal SUVR adjusted with the GCI in amnesic typical AD patients. We are not aware of any previous data evaluating the relationship between HSV-1 serology and molecular neuroimaging of Aß or Tau lesions in AD patients. Earlier reports, obtained in cohorts of elderly but non-demented patients, have so far shown contradictory results: positive relationships between IgG titers and Aß SUVR in frontal brain regions (Cantero et al., 2024), and conversely, a negative relationship (trend) between anti-HSV-1 antibody titers and cortical amyloid burden (Linard et al., 2025).

The association between HSV-1 infection and both Aß and Tau burden in the temporal lobe involvement is particularly relevant since it is known that the temporal lobes, the hippocampus, and the amygdala are the preferred targets of HSV-1 during acute infections such as herpes encephalitis (Damasio et al., 1985) and are also regions with an enrichment of viral material (HSV-1 genome) in the brains of elderly subjects (Jamieson et al., 1992; Marques et al., 2001). Other *in vivo* studies in humans showed indeed that brain metabolism in the temporal cortex is negatively impacted by high viral activity (Goldhardt et al., 2022) and high anti-HSV-1 IgG titers are also associated with hippocampal atrophy (Linard et al., 2021). Surprisingly, we did not find associations between HSV-1 seropositivity and regional index in the precuneus gyrus, another brain area presenting early ADNC (Levin et al., 2021). This could support the hypothesis of a preferential involvement of temporal regions in HSV-1-mediated neuropathological changes. The fact that we did not find any association between HSV-1 seropositivity and the regional Aß or Tau SUVR indices in atypical AD cases, for whom the limbic regions are more preserved (Whitwell et al., 2012; Whitwell et al., 2018), is consistent with this hypothesis. This interpretation should however be taken with caution. Indeed, it is impossible to rule out an absence of effects due to a lack of statistical power in the atypical subgroup regression analysis. It would be useful to confirm these results in larger clinical studies including information on viral reactivations and by studying the specificity of HSV-1 infection markers vs other peripheral infectious markers.

There are several additional limits to this work. In animal models, the exact nature of HSV-1 induced brainstem/cerebellum Aß deposits should be refined, in particular in terms of topographies and relationships with characterized inflammatory responses that were found to be spatially associated. Also, the impact of viral reactivations, a key factor in the modulation of post-infection AD-like phenotypes <u>(De Chiara et al., 2019)</u>, may be addressed in the cotton rat model. Our first attempts to induce viral reactivation in cotton rats (by thermal stress, UV or corticoids) were unsuccessful and deserve further efforts. Neuropathological postmortem data in human tissue should also be completed using an expanded list of brain areas and techniques like RNAscope ISH to better appreciate the topographies of HSV-1 genomes, their status (lytic/latent) and their association with Aß and Tau proteinopathies. The results obtained in the ShaTau7-Imatau subcohort are clearly limited by the low number of analyzed subjects. Larger samples would allow 1) an assessment of the effect of APOE4, an important modulator of the herpes-AD interplay (Urosevic et al., 2008), 2) evaluating anti-HSV-1 IgG levels or anti-HSV-1 IgM to take into account the critical impact of reactivation episodes (Linard et al., 2021; Linard et al., 2020; Lopatko Lindman et al., 2019) and 3) refine analysis in typical vs atypical AD variants.

Despite these limits, the results obtained in the cotton-rat model, illustrating *de novo* induction of Aβ and Tau pathology, after primary infection, in the brainstem/cerebellum and at a distance, as well as observations collected in humans (postmortem and in-vivo data), converge to support the hypothesis that HSV-1 is involved in initiating or modulating the pathophysiology of AD.

## Acknowledgments

Part of this work was funded by a grant from France Alzheimer to BD. The research leading to these results also received funding from the national program “Investissements d’avenir” ANR-10- IAIHU-0006.

The authors gratefully acknowledge the staff of Histomics and ICM.Quant core facilities of ICM for the help in carrying out the research.

We would also like to thank Dr. Elodie Martin from the Lubetzki-Stankoff team at the ICM and G. Dorothée’s team (Saint-Antoine Hospital, Paris) for their valuable assistance in setting up the biochemical assay protocols. Dr Oscar Haigh, Dr. P. Lomonte and Pr. M. Labetoulle are greatly acknowledged for discussing some aspects of this work.

The Shatau7/Imatau study was funded by French Ministry of Health grant (PHRC- 2013-0919), CEA, Fondation Recherche Alzheimer, Institut de Recherches Internationales Servier, Association France-Alzheimer.

We also thank the donors and the Brain Donation Program of the “The Brainbank Neuro-CEB Neuropathology Network” run by a consortium of Patient Associations with the support of Fondation Alzheimer and IHU A-ICM.

## Notes

### Competing Interest Statement

The authors have declared no competing interest.

